# Meiotic drive adaptive testes enlargement during early development in the stalk-eyed fly

**DOI:** 10.1101/2022.08.08.502295

**Authors:** Sasha L Bradshaw, Lara Meade, Jessica Tarlton-Weatherall, Andrew Pomiankowski

**Author notes:** **Author for correspondence:** Andrew Pomiankowski.

## Abstract

The sex-ratio ‘SR’ X-linked meiotic drive system in stalk-eyed flies destroys all Y-bearing sperm. Unlike other SR systems, drive males do not suffer fertility loss. They have greatly enlarged testes, which compensate for gamete killing. We predicted that enlarged testes arise from extended development with resources re-allocated from the accessory glands, as these tend to be smaller in drive males. To test this, we tracked the growth of the testes and accessory glands of wildtype and drive males over 5–6 weeks post-eclosion before males attained sexual maturity. Neither of the original predictions are supported by this data. Instead, we found that the drive-male testes were enlarged at eclosion, reflecting a greater allocation of resources to the testes during pupation. In addition, there was no evidence that the greater allocation of resources to the testes during adult development retarded accessory gland growth. There was evidence of a general trade-off with eyespan, as males with larger relative eyespan had larger accessory glands but smaller testes. These findings support the idea that enlarged testes in drive males arise as an adaptive allocation of resources to traits that enhance male reproductive success.

**One sentence summary:** Adaptive testes enlargement in early development ensures maintenance of fertility in stalk-eyed flies that lose half of their sperm due to meiotic drive

## 1. Introduction

Mendel’s first law of equal segregation holds for most nuclear genes. This fair segregation can be subverted by meiotic drivers that gain a transmission advantage, often in conflict with the rest of the genome (1–3). Male meiotic drivers are genetically and mechanistically diverse, but all result in the death or disabling of noncarrier sperm (4). Many examples exist, both of autosomal (e.g. *SD* in *Drosophila melanogaster* (5,6), *t* locus in mice (7,8)), and sex-linked origin (e.g. *SR* in *D. simulans* (9,10), and *Slx*/*Sly* in mice (11)). Due to the dysfunction of wildtype sperm, meiotic drive detrimentally impacts male fertility (12,13), with drive sperm typically being less effective under sperm competition (8,9,13). In addition, meiotic drive is often associated with viability reduction in both males and females (14). Sex chromosome drive is also associated with various costs through the distortion of the population sex ratio (15–18), which can potentially lead to local extinction (19–21).

In response to these costs, host nuclear genes have been selected to resist drive or counter its deleterious effects (22–24). A common response is the evolution of drive suppressors (24–26). A number of putative behavioural adaptations are known. In the *t*-haplotype system in mice, juvenile dispersal is enhanced in *t* heterozygotes which reduces the probability of lethal homozygosity (27). Another example is the theoretical prediction that females should mate polyandrously to decrease the success of drive-bearing sperm (28). In alignment with this, experimental populations of *D. pseudoobscura* exposed to a high frequency of the SR meiotic drive evolved increased female remating (29), although there is no evidence that variation in drive frequency is a major factor determining female mating rate in wild populations (29). It has also been suggested that mate choice might allow females to discriminate against drive-carrying males, either through the pleiotropic effects of drive or via signals of genetic quality where drive is associated with reduced viability (14,30). There are some tentative examples, such as the major histocompatibility complex linked with the *t* haplotype in mice (31), and reduced eyespan in male stalk-eyed flies where female preference favours longer eyespan (32). But again, there is no evidence that the presence of drive has led to the strengthening of mate preferences.

We investigated evidence for an adaptive response in reproductive organ size to X-linked drive (SR) in the Malaysian stalk-eyed fly, *Teleopsis dalmanni*. Previous work has shown that SR males deliver the same number of sperm per ejaculate (33) and do not suffer fertility loss compared to wildtype males (34). This reflects a massive increase in the size of their testes which are ∼26% larger than wildtype (34). This could be due to a resource trade-off with the accessory glands which are reduced in SR males (34). In order to test this hypothesis, the testes and accessory glands of SR and wildtype males were dissected and measured over a series of developmental timepoints from eclosion to beyond the point of sexual maturity (35–37) to determine interactions in the growth profiles of these reproductive organs.

## 2. Materials and Methods

A wildtype (ST) stock was collected in 2005 from the Ulu Gombak Valley, Peninsular Malaysia (by A. Pomiankowski and S. Cotton). Flies with the X^SR^ gene were collected in 2012 from the same location and since 2019 have been maintained as a homozygous SR stock (16,38). Experimental ST males (X^ST^/Y) were collected on egglays (petri dish with damp cotton and sweetcorn) from cages housing X^ST^/X^ST^ females and X^ST^/Y males. The egglays were incubated at 25ºC and the emerging flies were collected daily. Males were housed by emergence date and females were discarded. The same procedure was followed to collect SR males (X^SR^/Y) from cages housing homozygous X^SR^/X^SR^ females and X^ST^/Y males.

Two experiments were conducted following the same method. The first experiment performed dissections over a longer period from day 0 (eclosion) to day 56 (“long dataset”). Flies were dissected on days 0, 1, 4, 8, 12, 16, 20, 34, and 56. A follow-up experiment was carried out with more intense measurements from day 11 to day 25 (“short dataset”), with dissections on days 11, 13, 15, 17, 19, 21, 23, and 25. The thorax and eyespan of ice-anaesthetised flies were measured prior to dissection, using an Infinity Capture video microscope attached to a computer equipped with NIH image software (FIJI (Image J) Version 2.1.0/1.53c). The thorax was measured from the prothorax anterior tip, along the midline ending at the joint in-between the thorax and metathoracic legs (36). Eyespan was measured from the outer tips of the eyes adjacent to where the stalk joins the eye bulb (38). Flies were then dissected in 15ul of phosphate buffer saline (PBS) using 5mm forceps on a glass slide under the stereomicroscope. The testes and accessory glands were isolated then untangled and uncoiled without causing rupture or damage. Excess material such as external cuticle was removed from the slide to prevent distortion of the image. Another 15ul of PBS was added before adding a glass cover. Images were taken using a differential interference contrast microscope on QCapture Pro imaging software at either x5 or x10 magnification. The polygon selection tool in Image J was used to take area measurements for both testes and accessory glands, by tracing around the outline of the organs.

### Statistical Analysis

All statistical analyses were performed in R (Version 1.4.1103). Linear regression models were used to identify differences in reproductive organ size between genotypes. Models included genotype, age (days), thorax size, and residual eyespan. A stepwise build-up was used to add terms that improved the model fit. Terms that did not improve the model fit were discarded. The morphological traits of thorax size and relative eyespan were added as covariates as they are known to differ between genotypes, and correlate with reproductive organ size in mature adult flies (34). Relative eyespan was calculated from the residuals using a linear regression model after taking into account thorax size, as these traits are strongly collinear (39). To determine trade-offs between the development of the testes and accessory glands with other morphological traits, interaction terms were tested. Mean and standard error trait sizes (mm) are reported throughout. See supplementary information (SI) for all models.

## 3. Results

### Body size and eyespan

In the long dataset, the body size (thorax) of SR (mean ± s.e. = 2.324 ± 0.012) and ST males (2.352 ± 0.013) did not differ (F_1,367_ = 2.651, p = 0.104). The eyespan of SR was smaller than ST males (SR: 7.872 ± 0.056, ST: 8.095 ± 0.061, F_1,367_ = 7.266, p < 0.001), and this held after controlling for body size (F_1,366_ = 5.253, p < 0.010). In the short dataset, the body size of SR (2.441 ± 0.009) was smaller than ST males (2.511 ± 0.009; F_1,355_ = 29.327, p < 0.001). Once more, eyespan was smaller in SR (8.382 ± 0.034) than ST males (8.753 ± 0.032; F_1,355_ = 61.941, p < 0.001), and this held after controlling for body size (F_1,354_ = 31.197, p < 0.001).

### Testes and accessory glands

Controlling for the day of dissection, in the long data set, SR had larger testes (1.006 ± 0.050) than ST males (0.793 ± 0.040; F_1,356_ = 11.155, p < 0.001; Figure 1A, 2A) and this held after controlling for body size and relative eyespan (F_1,353_ = 57.2745, p < 0.001). The same was the case over the restricted timeframe of the short data set (SR: 1.202 ± 0.023, ST: 0.97 ± 0.018; F_1,344_ = 84.038, p < 0.001; Figure 1C, 2C), and again after controlling for body size and relative eyespan (F_1,342_ = 107.194, p < 0.001). Considering individual time points separately, SR male testes were larger on days 0, 1, 4, 8, 12, 20, and 56 (p < 0.05) but not on days 16 and 34 (p > 0.05; Table S1) in the long dataset. When repeated at the higher sample size in the short dataset, SR male testes were larger on days 11, 13, 17, 19, 21, 23, 25 (p < 0.05) but marginally not on day 15 (p = 0.052; Table S3).

**Figure 1.**
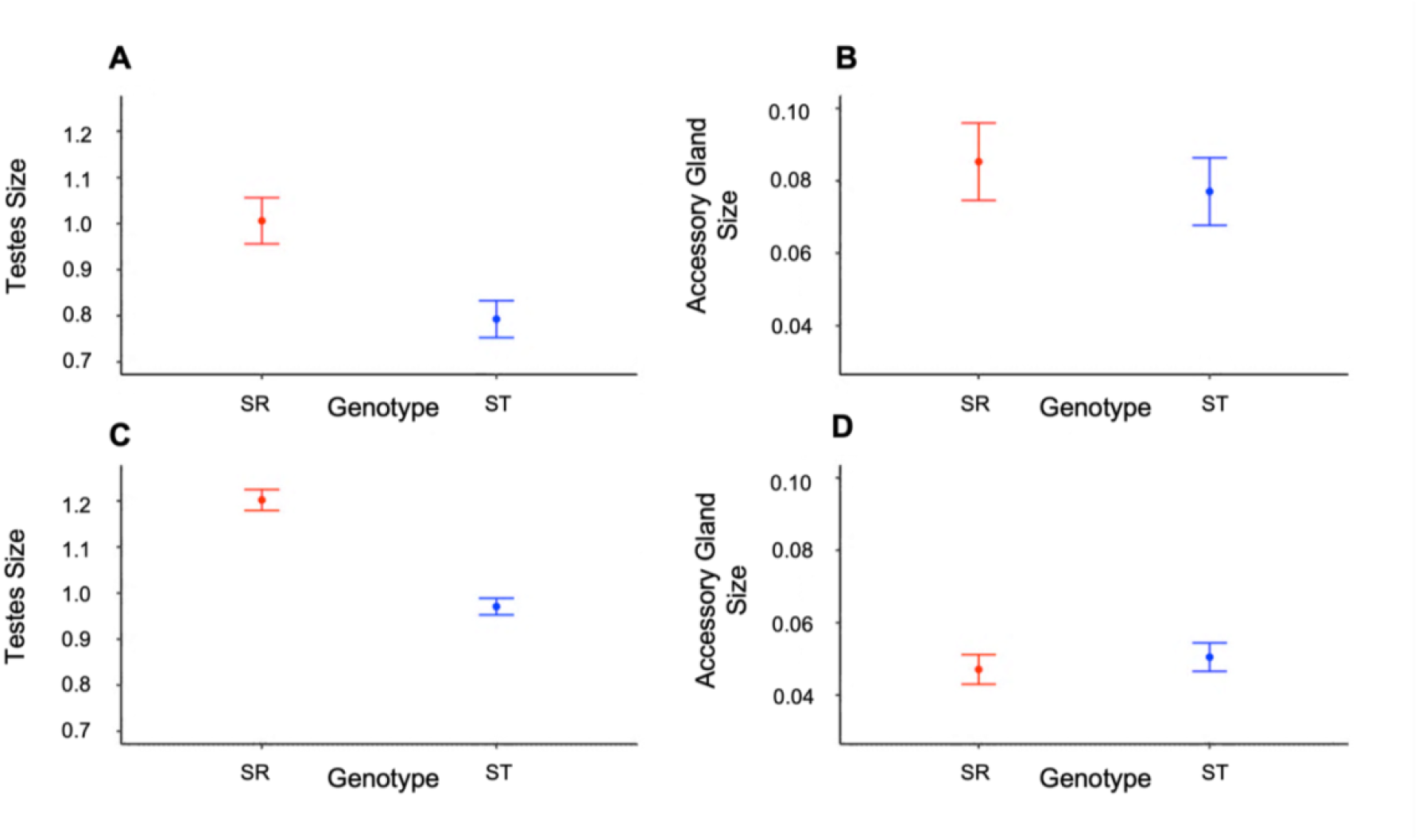
Size and growth of reproductive organs. Mean ± s.e. for SR (red) and ST (blue) testes size in the **(A)** long (0–56 days) and **(B)** short datasets (11–25 days), and similar measures for accessory gland size in the **(C)** long and **(D)** short datasets.

**Figure 2.**
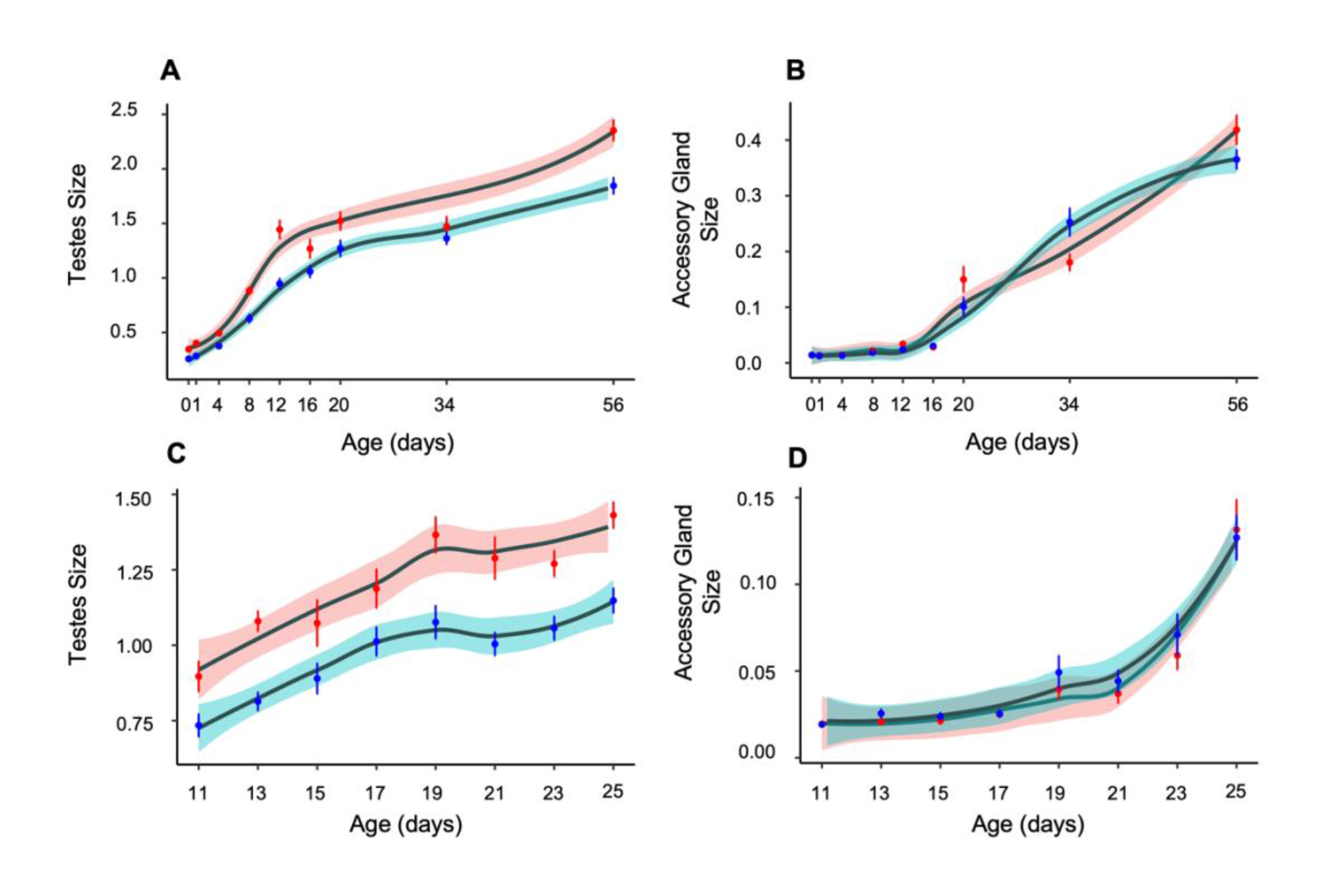
Growth curves across age (days since eclosion) for SR (red) and ST (blue) testes size in the **(A)** long and **(C)** short datasets, and for accessory gland size in the **(B)** long and **(D)** short datasets. Growth curves were fitted using the ‘loess’ function of the ‘ggplot2’ package in R (v1.4.1103); shaded area represents ± s.e.

Accessory gland growth contrasted with testes as there was no overall differences between SR and ST males in either the long (SR: 0.085 ± 0.012, ST: 0.077 ± 0.009, F_1,315_ = 0.339, p = 0.561; Figure 1B, 2B) or short dataset (SR: 0.047 ± 0.004, ST 0.051 ± 0.004, F_1,345_ = 0.358, p = 0.550; Figure 1D, 2D) or after controlling for body size and relative eyespan (Long: F_1,313_ = 0.425, p = 0.515, Short: F_1,343_ = 0.027, p = 0.870). There was likewise no difference on individual dissection days in either dataset (p > 0.05; Table S2, S4).

### Interactions between testes, accessory glands, and relative eyespan

The short dataset was investigated to identify whether there was a trade-off between the size of the reproductive organs during their most rapid period of growth. Males with larger accessory glands had larger testes after controlling for body size (F_1,338_ = 14.859, p < 0.001). The growth of the accessory glands was predicted by relative eyespan as males with larger relative eyespan had larger accessory glands (F_1,338_ = 10.965, p < 0.001). Testes size showed a more complex relationship as males with larger relative eyespan had larger accessory glands but smaller testes (F_1, 337_ = 6.021, p = 0.015; Figure S1). These relationships between relative eyespan and reproductive organ size did not differ with genotype.

## 4. Discussion

Male carriers of sex-ratio (SR) meiotic drive in *T. dalmanni* experience reduced sperm production due to the dysfunction of noncarrier sperm (33). They compensate for this with enlarged testes. The experiments here show that this reflects pupal resource allocation as SR testes size is already greater at eclosion (day 0). We found no evidence that greater SR testes size is a passive response to lower sperm production, as sperm bundles mature much later in development from 12 days post-eclosion (37). These findings suggest that the enlargement of the testes is an adaptation to compensate for the future loss of sperm caused by meiotic drive, allowing adult male sperm production and fertility to be maintained at the same level as wildtype males (34). This is likely to be encoded on the X^SR^ chromosome which contains a set of inversions spanning most of its length (40), which would allow tight linkage to be maintained with the genes responsible for controlling drive.

We previously hypothesized that the enlargement of the testes induced a resource trade-off with the accessory glands, which are smaller in SR adult males (34). There was no evidence for this, as greater testes size in SR males did not depress accessory gland size in either dataset, even though the second ‘short dataset’ (days 11–25) was designed to hone-in on the period when both reproductive organs undergo rapid development. There was also no evidence of a pupal resource allocation trade-off as accessory glands were the same size in SR and wildtype males at eclosion. In fact, there was no obvious difference in accessory gland size at any development stage from eclosion to day 56. This finding contrasts with previous experimental work (Meade et al. 2020) and data from the field (unpublished) showing reduced accessory gland size in SR males. A possible explanation is that flies used in the present study were virgins. In the study by Meade et al. (2020), dissections were performed on males, the majority of which had already mated with one or several females. Mating causes a decrease in accessory gland size (35,41), and we hypothesize that mated SR drive males may have a reduced capacity to replenish their accessory gland resources. There is indirect support for this idea, as SR males mate less when housed with multiple females, because they take longer to re-mate (34). This hypothesis will be addressed in a follow-up experiment.

Though there was no evidence of a resource trade-off between the size of the testes and accessory glands, we found evidence for a more complex trade-off involving the secondary sexual ornament of eyespan. Males with larger relative eyespan (after correction for body size) invest more in accessory gland size and less in testes size. These males are more attractive to females and have more mating opportunities, both in experimental work and in field samples (30,32,39,42). Larger accessory glands are likely to be beneficial to the individual, as their size strongly influences the re-mating frequency (41). For males with larger relative eyespan, investing in the accessory glands will give higher returns than investing in testes size. Conversely, males with smaller relative eyespan are less attractive, expect fewer matings, and gain less from investment in accessory glands. This relationship is apparent across genotypes, revealing that the same selective pressures even apply to SR males, which exhibit a general reduction in eyespan.

## Supporting information

Supplementary Statistics and Figures

## Acknowledgements

SB is supported by a Studentship from the London Natural Environment Research Council DTP (NE/S007229/1), AP is supported by funding from the Engineering and Physical Sciences Research Council (EP/F500351/1, EP/I017909/1), Natural Environment Research Council (NE/R010579/1) and Biotechnology and Biological Sciences Research Council (BB/V003542/1). We thank Flo Camus and Sadé Bates for help with statistical analyses in R, and Rebecca Finlay and Wendy Hart for help with maintenance of stalk-eyed fly stocks.

## Data and materials availability

Data will be submitted to the Dryad repository when the MS is accepted

