## Supplementary Statistics and Figures for "Meiotic drive adaptive testes enlargement during early development in the stalk-eyed fly"

**Supplementary material from "Meiotic drive-linked adaptive testes enlargement during early development in the stalk-eyed fly"**

**Authors**: Sasha Bradshaw, Lara Meade, Jess Weatherall and Andrew Pomiankowski

**Statistical Models and Tables S1-S4**

### Supplementary Information: Models

#### Traits (Long dataset)

**Body Size** The body size (thorax) of SR and ST males did not differ across all time points.

```
Anova(lm(thorax ~ genotype, data = long))
```

```
## Anova Table (Type II tests)
##
## Response: thorax
##            Sum Sq  Df F value Pr(>F)
## genotype   0.0747   1   2.651 0.1043
## Residuals 10.3389 367
```

**Eyespan** SR males had smaller eyespan than ST males, even after controlling for body size.

```
Anova(lm(eyespan ~ genotype, data = long))
```

```
## Anova Table (Type II tests)
##
## Response: eyespan
##            Sum Sq  Df F value   Pr(>F)
## genotype    4.569   1  7.2664 0.007349 **
## Residuals 230.740 367
## ---
## Signif. codes:  0 '***' 0.001 '**' 0.01 '*' 0.05 '.' 0.1 ' ' 1
```

```
Anova(lm(eyespan ~ genotype + thorax, data = long))
```

```
## Anova Table (Type II tests)
##
## Response: eyespan
##            Sum Sq  Df  F value  Pr(>F)
## genotype    1.234   1   5.2529 0.02248 *
## thorax    144.767   1 616.2950 < 2e-16 ***
## Residuals  85.973 366
## ---
## Signif. codes:  0 '***' 0.001 '**' 0.01 '*' 0.05 '.' 0.1 ' ' 1
```

**Relative Eyespan** Relative eyespan was calculated across all time points.

```r
resES <- residuals(lm(eyespan ~ thorax, data = long
$resES <- resES
```

**Testes** SR males had larger testes than ST males, even when controlling for age.

```r
Anova(lm(t_area ~ genotype, data = long))
```

```
## Anova Table (Type II tests)
##
## Response: t_area
##            Sum Sq  Df F value    Pr(>F)
## genotype    4.069   1  11.155 0.0009269 ***
## Residuals 129.866 356
## ---
## Signif. codes:  0 '***' 0.001 '**' 0.01 '*' 0.05 '.' 0.1 ' ' 1
```

```r
Anova(lm(t_area ~ genotype + age_days, data = long))
```

```
## Anova Table (Type II tests)
##
## Response: t_area
##           Sum Sq  Df F value    Pr(>F)
## genotype   4.773   1  49.626 9.713e-12 ***
## age_days  95.723   1 995.264 < 2.2e-16 ***
## Residuals 34.143 355
## ---
## Signif. codes:  0 '***' 0.001 '**' 0.01 '*' 0.05 '.' 0.1 ' ' 1
```

This held true after controlling for thorax and relative eyespan.

```r
Anova(lm(t_area ~ genotype + age_days + thorax + resES, data = long))
```

```
## Anova Table (Type II tests)
##
## Response: t_area
##           Sum Sq  Df  F value    Pr(>F)
## genotype   5.356   1  57.2745 3.331e-13 ***
## age_days  93.360   1 998.3266 < 2.2e-16 ***
## thorax     0.628   1   6.7113  0.009977 **
## resES      0.511   1   5.4649  0.019961 *
## Residuals 33.011 353
## ---
## Signif. codes:  0 '***' 0.001 '**' 0.01 '*' 0.05 '.' 0.1 ' ' 1
```

**Accessory Glands** There was no difference in accessory gland size between SR and ST males, even when controlling for age.

```r
Anova(lm(ag_area ~ genotype, data = long))
```

```
## Anova Table (Type II tests)
##
## Response: ag_area
##           Sum Sq  Df F value Pr(>F)
## genotype  0.0054   1   0.339 0.5608
## Residuals 4.9942 315
```

```
Anova(lm(ag_area ~ genotype + age_days, data = long))
```

```
## Anova Table (Type II tests)
##
## Response: ag_area
##           Sum Sq  Df   F value Pr(>F)
## genotype  0.0022   1    0.6968 0.4045
## age_days  4.0039   1 1269.5447 <2e-16 ***
## Residuals 0.9903 314
## ---
## Signif. codes:  0 '***' 0.001 '**' 0.01 '*' 0.05 '.' 0.1 ' ' 1
```

This held true after controlling for thorax and relative eyespan.

```
Anova(lm(ag_area ~ genotype + age_days + thorax + resES, data = long))
```

```
## Anova Table (Type II tests)
##
## Response: ag_area
##           Sum Sq  Df   F value Pr(>F)
## genotype  0.0029   1    0.9288 0.3359
## age_days  3.9018   1 1239.3589 <2e-16 ***
## thorax    0.0078   1    2.4675 0.1172
## resES     0.0004   1    0.1124 0.7377
## Residuals 0.9823 312
## ---
## Signif. codes:  0 '***' 0.001 '**' 0.01 '*' 0.05 '.' 0.1 ' ' 1
```

#### Traits (Short dataset)

**Body Size** The body size (thorax) of SR and ST males did not differ across all time points.

```
Anova(lm(thorax ~ genotype, data = short))
```

```
## Anova Table (Type II tests)
##
## Response: thorax
##           Sum Sq  Df F value    Pr(>F)
## genotype  0.4353   1  29.327 1.129e-07 ***
## Residuals 5.2693 355
## ---
## Signif. codes:  0 '***' 0.001 '**' 0.01 '*' 0.05 '.' 0.1 ' ' 1
```

**Eyespan** SR males had smaller eyespan than ST males, even after controlling for body size.

```
Anova(lm(eyespan ~ genotype, data = short))
```

```
## Anova Table (Type II tests)
##
## Response: eyespan
##           Sum Sq  Df F value    Pr(>F)
## genotype  12.275   1  61.941 4.326e-14 ***
## Residuals 70.350 355
## ---
## Signif. codes:  0 '***' 0.001 '**' 0.01 '*' 0.05 '.' 0.1 ' ' 1
```

```
Anova(lm(eyespan ~ genotype + thorax, data = short))
```

```
## Anova Table (Type II tests)
##
## Response: eyespan
##           Sum Sq  Df F value    Pr(>F)
## genotype   4.177   1  31.197 4.653e-08 ***
## thorax    22.957   1 171.479 < 2.2e-16 ***
## Residuals 47.392 354
## ---
## Signif. codes:  0 '***' 0.001 '**' 0.01 '*' 0.05 '.' 0.1 ' ' 1
```

**Relative Eyespan** Relative eyespan was calculated across all time points.

```
resES <- residuals(lm(eyespan ~ thorax, data = short
$resES <- resES
```

**Testes** SR males had larger testes than ST males, even when controlling for age.

```
Anova(lm(t_area ~ genotype, data = short))
```

```
## Anova Table (Type II tests)
##
## Response: t_area
##            Sum Sq  Df F value    Pr(>F)
## genotype   4.6355   1   63.17 2.722e-14 ***
## Residuals 25.3162 345
## ---
## Signif. codes:  0 '***' 0.001 '**' 0.01 '*' 0.05 '.' 0.1 ' ' 1
```

```
Anova(lm(t_area ~ genotype + age_days, data = short))
```

```
## Anova Table (Type II tests)
##
## Response: t_area
##            Sum Sq  Df F value    Pr(>F)
```

```
## genotype   4.5243   1  84.038 < 2.2e-16 ***
## age_days   6.7965   1 126.245 < 2.2e-16 ***
## Residuals 18.5196 344
## ---
## Signif. codes:  0 '***' 0.001 '**' 0.01 '*' 0.05 '.' 0.1 ' ' 1
```

This held true after controlling for thorax and relative eyespan.

```
Anova(lm(t_area ~ genotype + age_days + thorax + resES, data = short))
```

```
## Anova Table (Type II tests)
##
## Response: t_area
##            Sum Sq  Df  F value    Pr(>F)
## genotype   5.4652   1 107.1938 < 2.2e-16 ***
## age_days   7.3578   1 144.3154 < 2.2e-16 ***
## thorax     0.9420   1  18.4764 2.246e-05 ***
## resES      0.2147   1   4.2111   0.04092 *
## Residuals 17.4367 342
## ---
## Signif. codes:  0 '***' 0.001 '**' 0.01 '*' 0.05 '.' 0.1 ' ' 1
```

**Accessory Glands** There was no difference in accessory gland size between SR and ST males, even when controlling for age.

```
Anova(lm(ag_area ~ genotype, data = short))
```

```
## Anova Table (Type II tests)
##
## Response: ag_area
##            Sum Sq  Df F value Pr(>F)
## genotype  0.00100   1  0.3576 0.5503
## Residuals 0.96472 345
```

```
Anova(lm(ag_area ~ genotype + age_days, data = short))
```

```
## Anova Table (Type II tests)
##
## Response: ag_area
##            Sum Sq  Df  F value Pr(>F)
## genotype  0.00080   1   0.4377 0.5087
## age_days  0.33762   1 185.2017 <2e-16 ***
## Residuals 0.62710 344
## ---
## Signif. codes:  0 '***' 0.001 '**' 0.01 '*' 0.05 '.' 0.1 ' ' 1
```

This held true after controlling for thorax and relative eyespan.

```
Anova(lm(ag_area ~ genotype + age_days + thorax + resES, data = short))
```

```
## Anova Table (Type II tests)
##
## Response: ag_area
##            Sum Sq  Df  F value    Pr(>F)
## genotype  0.00024   1   0.1324  0.716205
## age_days  0.33320   1 187.0916 < 2.2e-16 ***
## thorax    0.00106   1   0.5940  0.441404
## resES     0.01755   1   9.8541  0.001842 **
## Residuals 0.60908 342
## ---
## Signif. codes:  0 '***' 0.001 '**' 0.01 '*' 0.05 '.' 0.1 ' ' 1
```

##### Tradeoff Interactions

Interaction terms were tested between the testes, accessory glands, and relative eyespan.

Males with larger accessory glands had larger testes after controlling for body size.

```
summary(lm(t_area ~ thorax + age_days + ag_area, data = short), type = "III")
```

```
##
## Call:
## lm(formula = t_area ~ thorax + age_days + ag_area, data = short)
##
## Residuals:
##      Min       1Q   Median       3Q      Max
## -0.55007 -0.19112 -0.02467  0.15282  0.90004
##
## Coefficients:
##             Estimate Std. Error t value Pr(>|t|)
## (Intercept) 0.351806   0.288033   1.221 0.222780
## thorax      0.115360   0.110379   1.045 0.296709
## age_days    0.021752   0.003665   5.935 7.28e-09 ***
## ag_area     1.180468   0.321794   3.668 0.000283 ***
## ---
## Signif. codes:  0 '***' 0.001 '**' 0.01 '*' 0.05 '.' 0.1 ' ' 1
##
## Residual standard error: 0.2535 on 339 degrees of freedom
##   (14 observations deleted due to missingness)
## Multiple R-squared:  0.2541, Adjusted R-squared:  0.2475
## F-statistic:  38.5 on 3 and 339 DF,  p-value: < 2.2e-16
```

```
Anova(lm(t_area ~ thorax + age_days + ag_area, data = short), type = "III")
```

```
## Anova Table (Type III tests)
##
## Response: t_area
##             Sum Sq  Df F value    Pr(>F)
## (Intercept) 0.0959   1  1.4918 0.2227799
## thorax      0.0702   1  1.0923 0.2967089
## age_days    2.2629   1 35.2191 7.281e-09 ***
## ag_area     0.8646   1 13.4571 0.0002833 ***
```

```
## Residuals   21.7812 339
## ---
## Signif. codes:  0 '***' 0.001 '**' 0.01 '*' 0.05 '.' 0.1 ' ' 1
```

Relative eyespan predicted accessory gland size, as males with larger relative eyespan had larger accessory glands.

```
summary(lm(ag_area ~ thorax + age_days + t_area + resES, data = short), type = "III")
```

```
##
## Call:
## lm(formula = ag_area ~ thorax + age_days + t_area + resES, data = short)
##
## Residuals:
##       Min        1Q    Median        3Q       Max
## -0.069144 -0.023925 -0.004971  0.013555  0.244417
##
## Coefficients:
##               Estimate Std. Error t value Pr(>|t|)
## (Intercept) -0.1164853  0.0466823  -2.495 0.013062 *
## thorax       0.0098283  0.0180295   0.545 0.586029
## age_days     0.0057448  0.0005462  10.518  < 2e-16 ***
## t_area       0.0335262  0.0086974   3.855 0.000139 ***
## resES        0.0200752  0.0060627   3.311 0.001029 **
## ---
## Signif. codes:  0 '***' 0.001 '**' 0.01 '*' 0.05 '.' 0.1 ' ' 1
##
## Residual standard error: 0.04135 on 338 degrees of freedom
##   (14 observations deleted due to missingness)
## Multiple R-squared:  0.3998, Adjusted R-squared:  0.3927
## F-statistic: 56.29 on 4 and 338 DF,  p-value: < 2.2e-16
```

```
Anova(lm(ag_area ~ thorax + age_days + t_area + resES, data = short), type = "III")
```

```
## Anova Table (Type III tests)
##
## Response: ag_area
##              Sum Sq  Df  F value    Pr(>F)
## (Intercept) 0.01065   1   6.2264 0.0130623 *
## thorax      0.00051   1   0.2972 0.5860294
## age_days    0.18920   1 110.6325 < 2.2e-16 ***
## t_area      0.02541   1  14.8589 0.0001387 ***
## resES       0.01875   1  10.9646 0.0010290 **
## Residuals   0.57804 338
## ---
## Signif. codes:  0 '***' 0.001 '**' 0.01 '*' 0.05 '.' 0.1 ' ' 1
```

When testes size was controlled for, males with larger relative eyespan had larger accessory glands but smaller testes.

```r
summary(lm(t_area ~ thorax + age_days + resES + ag_area + resES:ag_area, data = short), type = "III")
```

```
##
## Call:
## lm(formula = t_area ~ thorax + age_days + resES + ag_area + resES:ag_area,
##     data = short)
##
## Residuals:
##      Min       1Q   Median       3Q      Max
## -0.52786 -0.18346 -0.02211  0.15381  0.90488
##
## Coefficients:
##                Estimate Std. Error t value Pr(>|t|)
## (Intercept)    0.331169   0.285817   1.159   0.2474
## thorax         0.123271   0.109491   1.126   0.2610
## age_days       0.021407   0.003646   5.872 1.03e-08 ***
## resES          0.052294   0.056756   0.921   0.3575
## ag_area        1.453749   0.333335   4.361 1.72e-05 ***
## resES:ag_area -2.127056   0.866867  -2.454   0.0146 *
## ---
## Signif. codes:  0 '***' 0.001 '**' 0.01 '*' 0.05 '.' 0.1 ' ' 1
##
## Residual standard error: 0.2513 on 337 degrees of freedom
##   (14 observations deleted due to missingness)
## Multiple R-squared:  0.2714, Adjusted R-squared:  0.2606
## F-statistic: 25.11 on 5 and 337 DF,  p-value: < 2.2e-16
```

```r
Anova(lm(t_area ~ thorax + age_days + resES + ag_area + resES:ag_area, data = short), type = "III")
```

```
## Anova Table (Type III tests)
##
## Response: t_area
##                Sum Sq  Df F value    Pr(>F)
## (Intercept)    0.0848   1  1.3425   0.24741
## thorax         0.0800   1  1.2675   0.26103
## age_days       2.1766   1 34.4761 1.034e-08 ***
## resES          0.0536   1  0.8489   0.35751
## ag_area        1.2008   1 19.0203 1.721e-05 ***
## resES:ag_area  0.3801   1  6.0208   0.01464 *
## Residuals     21.2756 337
## ---
## Signif. codes:  0 '***' 0.001 '**' 0.01 '*' 0.05 '.' 0.1 ' ' 1
```

**Table S1:** Welch two sample t-tests comparing **testes size** between genotypes on each day of the **long data set**

| Day | Mean SR | Mean ST | df | t | p-value |
|---|---|---|---|---|---|
| 0 | 0.345 | 0.258 | 42.197 | 3.333 | **0.002** |
| 1 | 0.400 | 0.284 | 42.399 | 4.637 | **<0.0001** |
| 4 | 0.495 | 0.375 | 43.899 | 3.865 | **<0.001** |
| 8 | 0.882 | 0.628 | 40.329 | 4.362 | **<0.0001** |
| 12 | 1.444 | 0.947 | 28.799 | 4.816 | **<0.0001** |
| 16 | 1.268 | 1.058 | 25.811 | 1.931 | 0.065 |
| 20 | 1.522 | 1.271 | 33.298 | 2.144 | **0.039** |
| 34 | 1.473 | 1.362 | 16.270 | 1.009 | 0.328 |
| 56 | 2.353 | 1.845 | 26.107 | 3.984 | **<0.001** |

**Table S2:** Welch two sample t-tests comparing **accessory gland size** between genotypes on each day of the **long data set**

| Day | Mean SR | Mean ST | df | t | p-value |
|---|---|---|---|---|---|
| 0 | 0.014 | 0.014 | 33.210 | 0.099 | 0.922 |
| 1 | 0.014 | 0.013 | 33.145 | 0.786 | 0.438 |
| 4 | 0.015 | 0.013 | 40.584 | 1.333 | 0.190 |
| 8 | 0.022 | 0.019 | 32.411 | 1.993 | 0.055 |
| 12 | 0.034 | 0.024 | 31.791 | 2.289 | **0.029** |
| 16 | 0.029 | 0.030 | 28.350 | -0.558 | 0.581 |
| 20 | 0.150 | 0.101 | 27.465 | 1.648 | 0.111 |
| 34 | 0.180 | 0.253 | 24.114 | -2.395 | **0.025** |
| 56 | 0.419 | 0.365 | 24.651 | 1.630 | 0.116 |

**Table S3:** Welch two-sample t-tests comparing **testes size** between genotypes on each day of the **short data set**

| Day | Mean SR | Mean ST | df | t | p-value |
|---|---|---|---|---|---|
| 11 | 0.897 | 0.734 | 38.106 | 2.610 | **0.013** |
| 13 | 1.080 | 0.814 | 42.690 | 5.981 | **<0.0001** |
| 15 | 1.073 | 0.890 | 36.095 | 2.013 | 0.052 |
| 17 | 1.188 | 1.012 | 34.689 | 2.205 | **0.034** |
| 19 | 1.366 | 1.076 | 35.762 | 3.626 | **<0.001** |
| 21 | 1.289 | 1.004 | 27.942 | 3.589 | **0.001** |
| 23 | 1.270 | 1.056 | 41.874 | 3.727 | **<0.001** |
| 25 | 1.431 | 1.148 | 50.93 | 4.783 | **<0.0001** |

**Table S4:** Welch two-sample t-tests comparing **accessory gland size** between genotypes on each day of the **short data set**

| Day | Mean SR | Mean ST | df | t | p-value |
|---|---|---|---|---|---|
| 11 | 0.020 | 0.019 | 37.478 | 0.139 | 0.890 |
| 13 | 0.021 | 0.026 | 28.923 | -1.773 | 0.087 |
| 15 | 0.021 | 0.024 | 40.221 | -0.889 | 0.379 |
| 17 | 0.025 | 0.025 | 41.506 | -0.166 | 0.869 |
| 19 | 0.040 | 0.049 | 28.765 | -0.868 | 0.393 |
| 21 | 0.037 | 0.044 | 36.521 | -0.884 | 0.383 |
| 23 | 0.059 | 0.071 | 35.263 | -0.846 | 0.403 |
| 25 | 0.132 | 0.127 | 47.485 | 0.213 | 0.832 |

**Supplementary material from "Meiotic drive-linked adaptive testes enlargement during early development in the stalk-eyed fly"**

**Authors**: Sasha Bradshaw, Lara Meade, Jess Weatherall and Andrew Pomiankowski

**Figure S1**

#### Figure S1

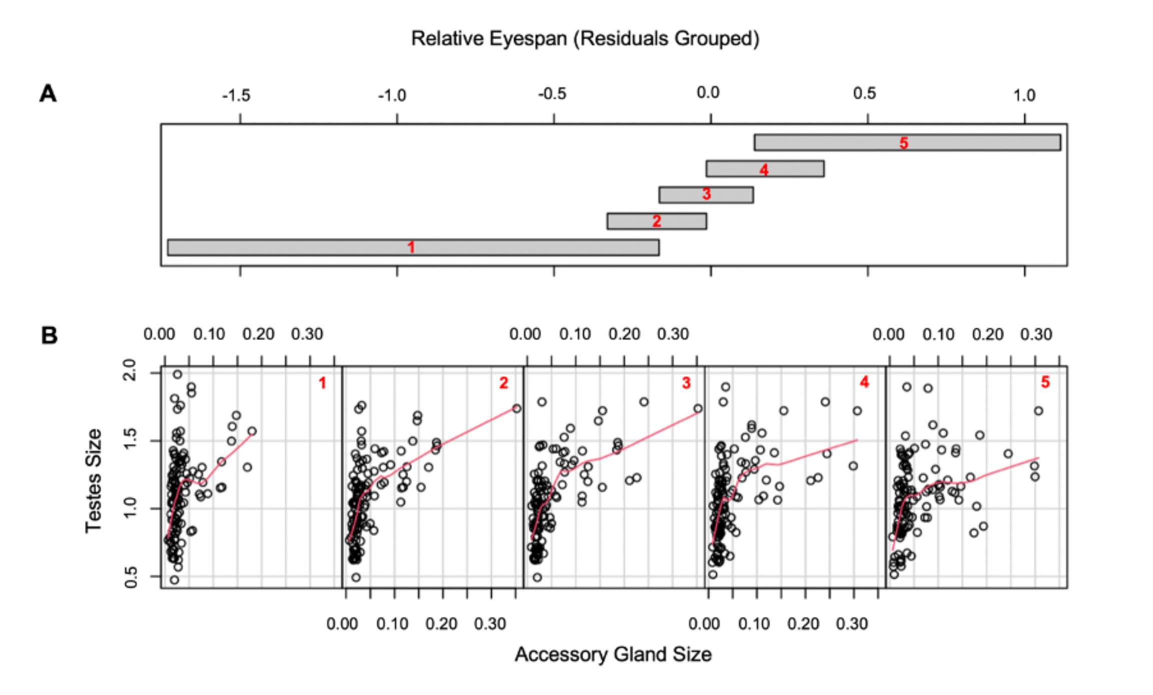


**Figure S1:** Interpretation of the trade-off between relative eyespan, accessory gland size and testes size. The 3-way interaction between these variables was investigated using conditioning plots using R-burble. For these plots, (**A**) the dataset is split up into a number of overlapping equal-sized regions (1-5) defined by relative eyespan (the conditioning variable). (**B**) Plots of the 5 regions show that as relative eyespan increases, males had larger accessory glands but smaller testes.
